# Historical Contingency Shapes Toxin Resistance in a Specialist Avian Predator

**DOI:** 10.1101/2025.07.04.662692

**Authors:** S. Mohammadi, S. Pradhan, F.G. Hoffmann, S. Herrera-Álvarez, Y. Deng, A. Eacock, S. Dobler, J. F. Storz, H.M. Rowland

## Abstract

Adaptations to toxic diets can cascade through ecosystems, altering physiology, species interactions, and trophic dynamics. To uncover the molecular basis of toxin resistance in the black-headed grosbeak (*Pheucticus melanocephalus*), a specialist predator of cardiotonic steroid-defended monarch butterflies (*Danaus plexippus*), we investigated the evolution of target-site insensitivity in the toxin’s molecular target – the Na,K-ATPase. Functional assays of engineered Na,K-ATPases revealed that resistance in grosbeaks arises from a non-additive interaction between a substitution at position 111 (Q111E) and up to six nearby amino acid changes in the first extracellular loop of the protein. Using resurrected ancestral proteins, we show that the earliest of these six substitutions to evolve (V113L) altered the functional effects of Q111E, such that Q111E alone became maladaptive. Only after the accumulation of additional permissive substitutions could Q111E confer resistance, highlighting how intramolecular epistatic interactions and historical contingency constrained the evolutionary path to adaptation. Our phylogenetic analysis of Na,K-ATPase sequences from 360 birds further revealed that several of the specific grosbeak substitutions—particularly at sites 112, 114, and 116—show strong signatures of co-evolution with changes at site 111 across the avian tree, supporting the hypothesis that resistance evolves through repeated, interacting changes. Together, these results reveal the molecular mechanisms of convergent evolution of toxin resistance and demonstrate how genetic background can shape evolutionary outcomes across trophic levels.

## Introduction

Cardiotonic steroids (CTSs) are a diverse class of toxins found in plants, insects, and vertebrates [1, 2]. They are highly toxic to most animals because they bind to and inhibit the α subunit of the Na,K-ATPase (ATP1A1), a transmembrane ion pump that is essential for maintaining membrane potentials and cellular function [3, 4]. The black-headed grosbeak (*Pheucticus melanocephalus*), a migratory songbird that breeds in western North America and winters in Mexico [5], exhibits a rare and highly specialised predatory behaviour. These birds prey on overwintering adult monarch butterflies (*Danaus plexippus*) in the Oyamel fir forests of central Mexico [6]. These butterflies are defended by CTS toxins [6, 7]. In mixed-species flocks with black-backed orioles (*Icterus galbula abeillei*), black-headed grosbeaks can kill up to 15,000 monarchs per day across the overwintering sites [8]. Analyses of grosbeak stomach contents have found CTS quantities as high as 259ug (equivalent to 6.17 mg/kg; [7]), and the birds can tolerate oral doses of 1050 mg/kg [6] – a level exceeding the lethal dose (LD_50_) of other CTSs tested in mice [9-13]. This remarkable resistance to CTSs has been suggested to be a further example of convergent molecular evolution in target-site insensitivity in the Na,K-ATPase [14, 15], a pattern documented in other vertebrates such as murid rodents, neotropical grass frogs, and snakes that feed on CTS-defended prey [16-18].

The molecular basis of CTS resistance has repeatedly evolved through amino acid substitutions in the first extracellular loop of Na,K-ATPase, which forms a critical region of the toxin’s binding pocket [16, 18-22]. Following this pattern, the black-headed grosbeak has a glutamine-to-glutamic acid substitution in the same loop at position 111 (Q111E). This amino acid site has been identified as a hotspot for major-effect substitutions that confer target-site insensitivity of Na,K-ATPase to CTS in several insect and vertebrate species [23, 24]. In addition to Q111E, there are five other amino acid changes in the grosbeak’s first extracellular loop – S112T, M114V, E115N, E116G, and P118G. This differentiates the grosbeak from its closest relative with an available genome sequence the Lazuli bunting (*Passerina amoena*) and from other species of birds that do not prey on monarchs (Fig. 1). L113, another substitution in the first extracellular loop distinguishes the grosbeak and other songbirds (Passeriformes) from most other species of birds, except for the Abyssinian ground hornbill (*Bucorvus abyssinicus*), which is also reported to feed on CTS-defended prey [25]. If these substitutions are responsible for the grosbeak’s ability to specialise on monarchs, the adaptation would represent a rare example of an evolutionary cascade [15, 26], whereby plant-derived toxins in monarchs drive selective pressures that extend to the third trophic level, shaping species far removed from the primary toxin source and, ultimately, species interactions across trophic levels.

**Figure 1.**
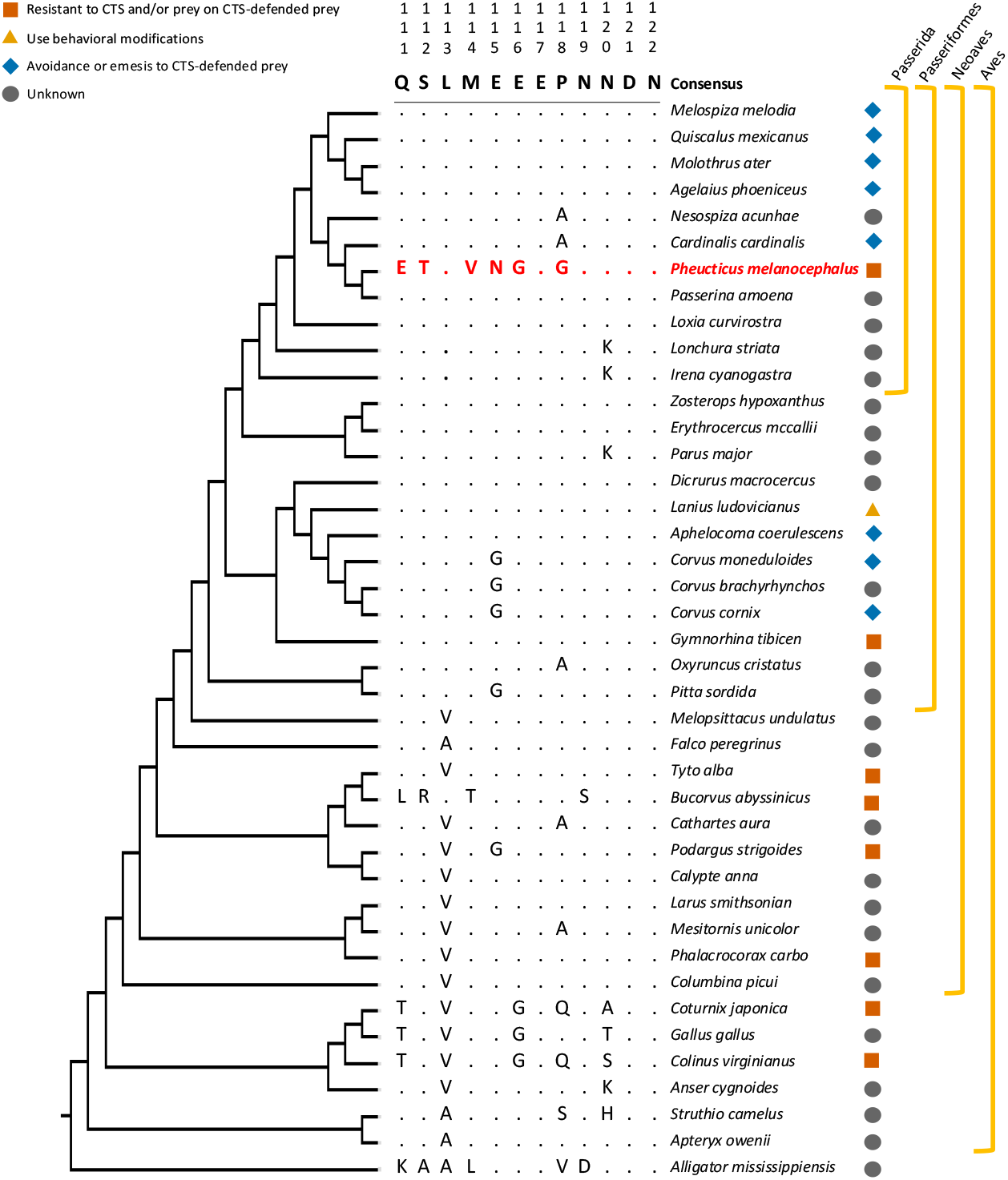
A phylogenetic subtree of birds showing amino acid differences in the first extracellular loop of the Na,K-ATPase α1 subunit (ATP1A1), a key target of CTSs. Amino acid positions are numbered according to the sheep (*Ovis aries*) reference sequence, following convention. Differences from the consensus sequence (not the inferred ancestral sequence) are shown for each species. Coloured symbols to the right of each species indicate the reported response of each species to CTS-defended prey: blue diamond – species that exhibit avoidance or emesis (vomiting) in response to CTS-defended prey; yellow triangle – species that use behavioural adaptations to feed on CTS-defended prey; orange square – species that show no emetic response when fed CTS-defended prey and/or have been observed feeding on CTS-defended prey or plants; grey circle – species for which no published data are available regarding CTS sensitivity or tolerance.

## Results and Discussion

We used protein engineering experiments to address the following questions about the grosbeak’s Na,K-ATPase. (1) Is Q111E a major-effect substitution that is necessary and sufficient to confer target-site insensitivity? (2) Does the set of associated substitutions (T112A, L113A, V114M, N115E, G116E, and G118P; hereafter referred to as 6subs) also contribute to target-site insensitivity? (3) do these additional substitutions compensate for deleterious effects of Q111E on overall protein function, rather than directly contributing to target-site insensitivity? To address these questions, we used heterologous protein expression and biochemical assays to quantify the functional effects of engineered variants in the Na,K-ATPase. Using the wildtype grosbeak Na,K-ATPase as a starting point, we synthesised four Na,K-ATPase constructs in which the Q111E and 6subs substitutions were individually or jointly reverted to their inferred ancestral state. For each engineered protein we measured resistance as the concentration of CTS (ouabain) at which enzyme activity is reduced to 50% (i.e., IC50) and assessed overall protein activity by quantifying the rate at which the enzyme hydrolyses ATP.

Consistent with the hypothesis that Q111E is a major-effect substitution with respect to target-site insensitivity, reverting that site to the ancestral amino acid state (E111Q) significantly reduced resistance in wildtype grosbeak Na,K-ATPase. We also found that reverting the amino acid states of 6subs significantly reduced resistance compared to the wildtype protein, though this reduction was significantly less pronounced than the effect of E111Q alone. Reverting Q111E and 6subs in tandem did not produce any further reduction in resistance relative to that produced by either reversion alone (Table S2, Fig. 2A). While our results clearly demonstrate the presence of target-site insensitivity in the grosbeak Na,K-ATPase, they also reveal that the resistance-conferring effects of E111 are conditional on the presence of additional amino acid substitutions within the extracellular loop.

**Figure 2.**
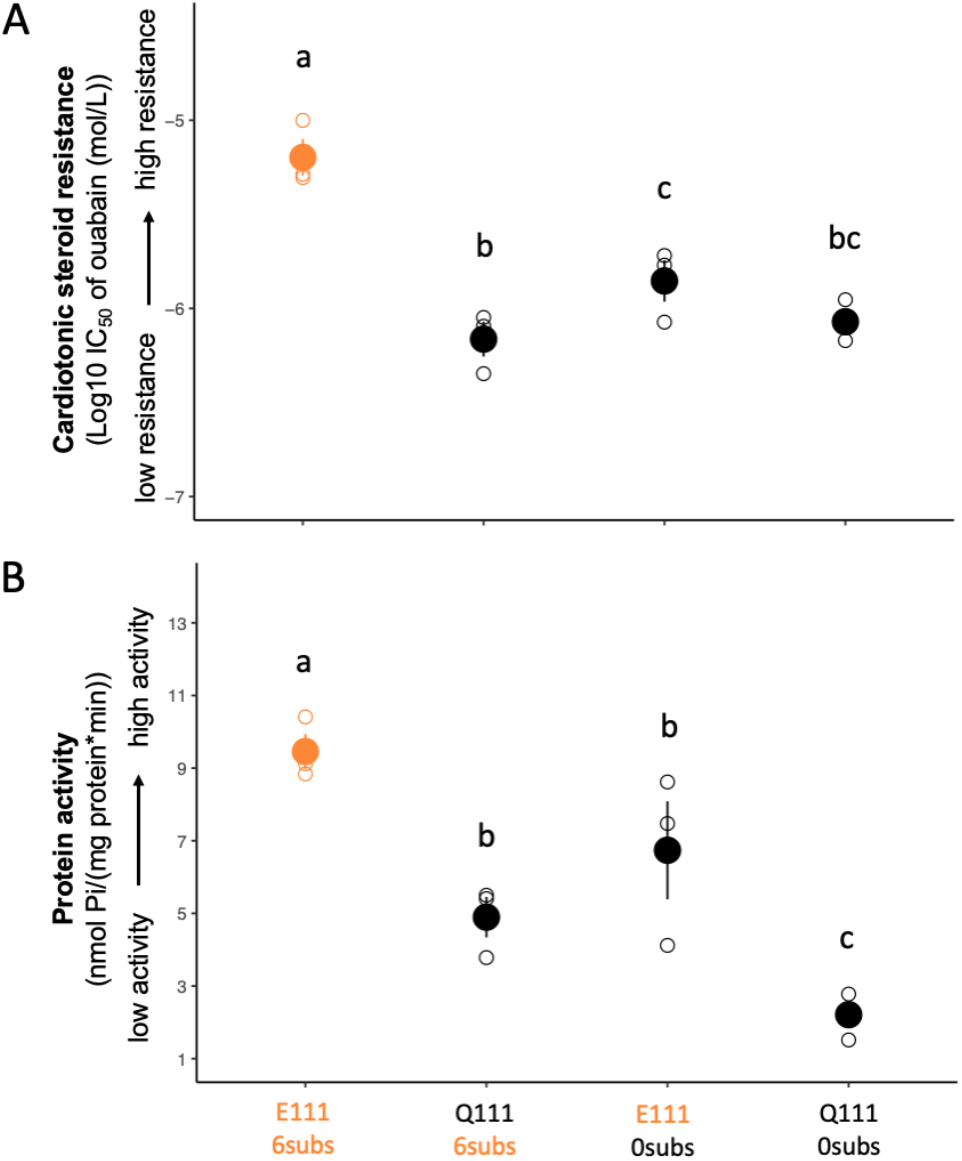
Functional effects of reverting grosbeak-specific Na,K-ATPase substitutions into their ancestral **s**tates, individually and in combination. For each recombinant protein, the substitutions were generated on the modern grosbeak genetic background and the resulting amino acid states are denoted on the x-axis. Amino acid states that are wildtype on the modern grosbeak Na,K-ATPase are given in orange and those that are ancestral in black. Panel A shows cardiotonic steroid resistance (i.e., mean log_10_IC_50_ ± SEM) plotted on the y-axis. Panel B shows the corresponding protein activity (i.e., mean ATP hydrolysis rate ± SEM) for the same proteins plotted on the y-axis. Raw data from three biological replicates are shown as open circles. Significant differences between proteins (contrast analysis) are indicated by different letters.

That reverting Q111E and the associated 6subs together does not reduce resistance beyond either reversion alone indicates a non-additive interaction, likely due to intramolecular epistasis. Similar nonadditive effects of resistance-conferring mutations have been documented in other systems of target-site insensitivity to CTS [16, 18]. For example, introducing E111 into the *Drosophila* Na,K-ATPase conferred only a tenfold increase in resistance, whereas the addition of a second substitution, Y122 (as found in aphids), led to more than a hundredfold increase in resistance [27]. In contrast to most previously studied systems, where epistasis typically involves two key sites (primarily positions 111 and 122), the grosbeak Na,K-ATPase shows conditional effects involving up to six interacting substitutions. This represents a more complex and unique evolutionary trajectory within a system otherwise characterised by highly repeatable molecular adaptations [28].

Functional investigations of CTS resistance-conferring substitutions in the Na,K-ATPase of other resistant species have revealed they often have negative pleiotropic effects on native activity (the rate at which the enzyme hydrolyses ATP), and such effects are sometimes mitigated by compensatory substitutions at other sites in the proteins [16, 18, 23, 24]. If the amino acid changes that confer CTS resistance in the grosbeak Na,K-ATPase negatively impact the protein’s other functions, we would predict that the individual reversions of Q111E and 6subs would produce a significant increase in protein activity relative to the wildtype grosbeak ATP1A1 genotype. Our experimental results show that this is not the case. Reverting Q111E and 6subs individually and in combination reduced protein activity (Fig. 2B; Table S2). The increased resistance to CTS in the grosbeak Na,K-ATPase is therefore associated with increased protein activity, which suggests that the mutation(s) conferring resistance also enhance or stabilise enzyme efficiency. In contrast to the compensatory amino acid interactions documented in the Na,K-ATPases of other CTS-resistant taxa [16, 18, 23, 24, 29], our experiments on grosbeak Na,K-ATPase provide no evidence that the resistance conferred by Q111E compromises native activity [16, 18, 23, 24, 29].

That E111’s resistance-conferring effects depend on the presence of multiple other substitutions indicates that the path to target-site insensitivity was constrained by earlier molecular changes. To determine whether historical contingency played a role in shaping the evolution of CTS resistance in grosbeaks, we statistically reconstructed and experimentally resurrected a set of progressively more ancient Na,K-ATPase constructs representing a temporal sequence of ancestral genotypes from three points in the grosbeak line of descent. These three points represent (1) the common ancestor of all birds (AncAves, 150–165 Mya [30] (Fig. 3A); (2) the common ancestor of Neoaves (AncNeoaves, 67.4 Mya [31]), an incredibly diverse groups of modern birds, including waterfowl (Anseriformes), shorebirds (Charadriiformes), raptors (Accipitriformes), and songbirds (Passeriformes); and (3) the common ancestor of Passerida (AncPasserida, 22.4 Mya [31]), a large and diverse clade (or infraorder) within the order Passeriformes, which includes finches, warblers, thrushes, sparrows, swallows, and larks. We selected AncPasserida as the most recent common ancestor of modern grosbeaks and other core passerines, rather than the common ancestor with *Passerina amoena* (the grosbeak’s closest sequenced relative), based on the hypothesis that the grosbeak’s specialised foraging behaviour on CTS-defended prey is an older rather than a more recent adaptation [32]. AncPasserida also provides a more conservative baseline for identifying derived substitutions in the grosbeak lineage (Table S3).

**Figure 3.**
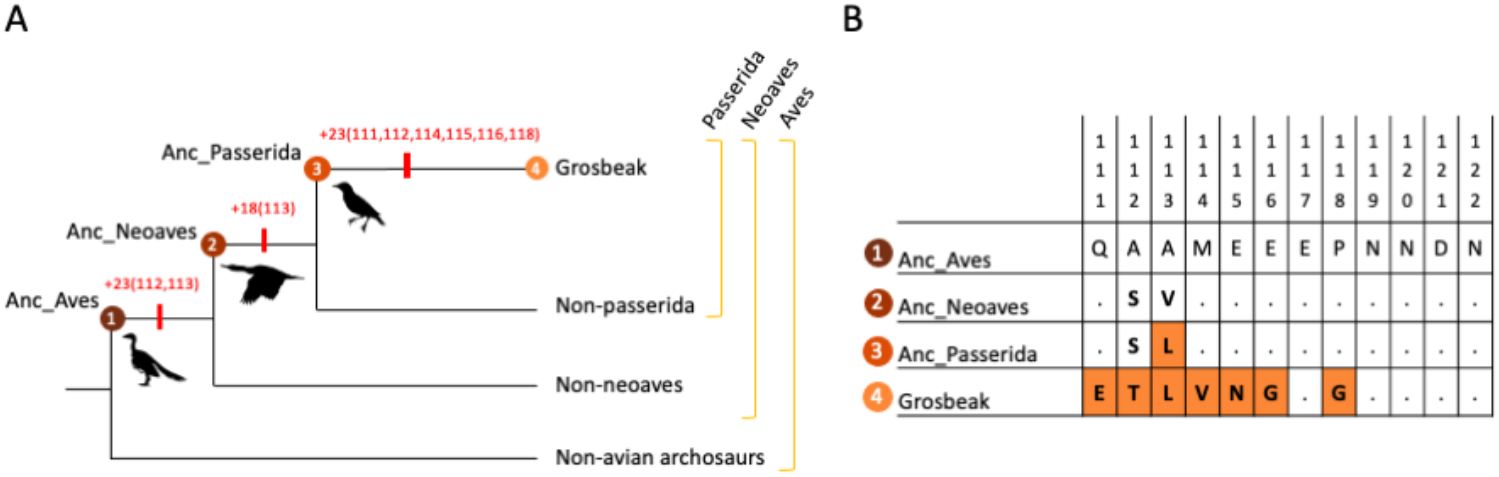
(A) A schematic phylogeny tracing the line of descent of the black-headed grosbeak back to the most recent common ancestor of all birds (AncAves). We reconstructed three ancestral steps along this lineage: ancestral Aves (node 1), ancestral Neoaves (node 2), and ancestral Passerida (node 3). From AncAves to the modern grosbeak, a total of 47 amino acid substitutions were gained in the Na,K-ATPase α1-subunit (encoded by the ATP1A1 gene). Of these substitutions, 23 (including A112S and A113V in the first extracellular loop) occurred on the branch connecting AncAves to AncNeoaves, 18 occurred on the branch connecting AncNeoaves to AncPasserida (including V113L), and 23 occurred on the branch connecting AncPasserida to the ATP1A1 of contemporary grosbeaks (including Q111E and the five remaining changes comprising 6subs; Fig. 3B). (B) Seven of these 47 substitutions (highlighted in orange) are in the protein’s first extracellular loop, which spans amino acid sites 111 to 122 and makes up a critical region of the toxin’s binding pocket.

We assessed whether the resistance-conferring substitutions found in modern grosbeaks would have had similar phenotypic effects in progressively more ancient ATP1A1 genotypes along their evolutionary lineage. To do this, we synthesised and heterologously expressed AncPasserida, AncNeoaves, and AncAves with and without the modern grosbeak substitutions Q111E and 6subs. By comparing the effects of Q111E and 6subs on these ancestral backgrounds, each progressively more divergent from the wildtype ATP1A1 genotype of contemporary grosbeaks, we tested how genetic context modulates the functional impact of these mutations

We found that toxin resistance conferred by Q111E alone was ancestrally accessible, but its effect diminished and eventually reversed as the mutation was moved onto increasingly more recent ancestral backgrounds (Fig. 4A; Table S4). Q111E caused a significant increase in resistance on the AncAves protein (Fig. 4A, left panel; Table S4). Q111E’s positive effect on resistance decreased when the mutation was introduced into the more recent AncNeoaves protein, which differs from AncAves by two substitutions (A112S and A113V) in the first extracellular loop. Subsequently, with the emergence of V113L in AncPasserida – the most recent ancestral background – the effect of Q111E reversed, shifting from resistance-increasing to resistance-decreasing (Fig. 4A, left panel; Table S4).

**Figure 4.**
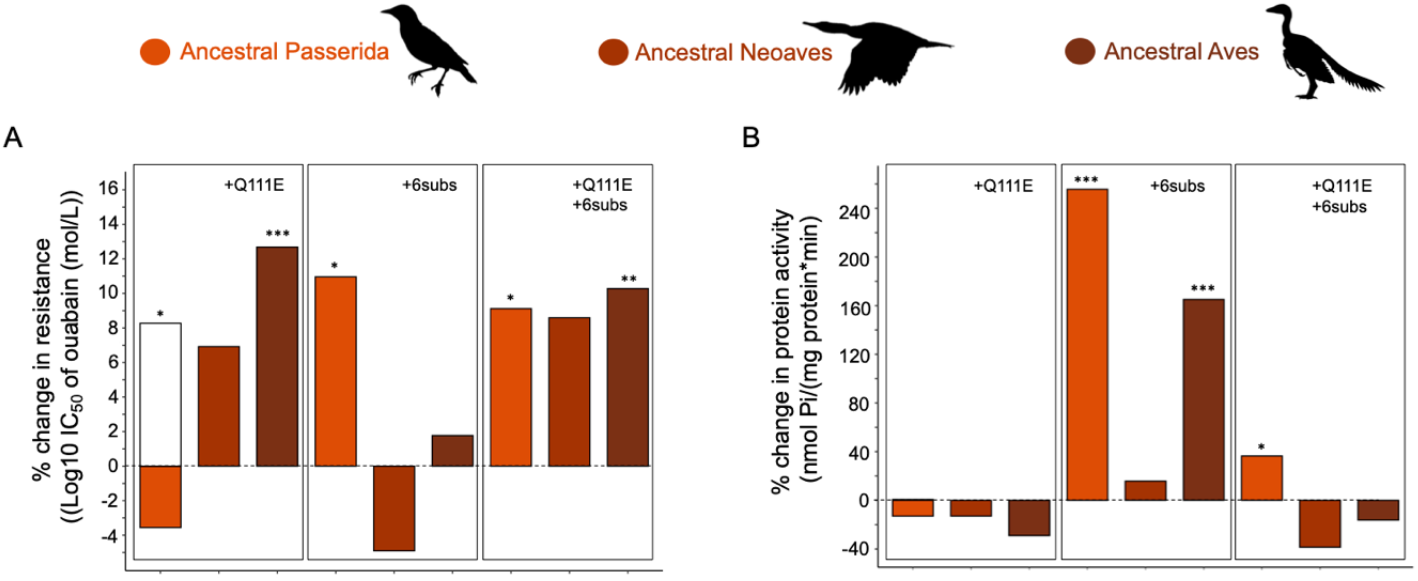
Functional properties of engineered Na,K-ATPases measuring the effects of modern black-headed grosbeak substitutions on increasingly more ancient genetic backgrounds (from orange to maroon). The graphs in panels A and B show the percent change in functional properties of wildtype ancestral NKAs in comparison to their mutagenised counterpart. Each set of bars (from left to right) shows the effects of one of three substitutions or sets of substitutions (Q111E, 6subs, and Q111E+6subs). In panel A, a measure of CTS resistance (i.e., mean log_10_IC_50_ ± SEM from three biological replicates) is plotted on the y axis. In panel B, a measure of protein activity (i.e., mean ATP hydrolysis rate ± SEM from three biological replicates) for the same proteins is plotted on the y axis. Significant differences (contrast analysis; Table S5, Fig. S1) between wildtype and mutagenised proteins are indicated by asterisks. In the Q111E graphs are an additional mutagenised construct of ancestral Passerida, engineered to carry Q111E as well as L113V. This additional construct is illustrated by a white bar. In panel B, this white bar completely overlaps with the original and is therefore not visible. Significance codes: 0 ‘***’; 0.001 ‘**’; 0.01 ‘*’.

Because V113L was the first substitution of the grosbeak’s 6subs to arise in AncPasserida, we investigated its potential role in altering the effect of Q111E. To do this, we synthesised and expressed an alternative AncPasserida, engineered to carry Q111E along with a reversion of V113L (i.e., L113V; a new version of AncNeoaves with 17 additional substitutions outside of the first extracellular loop; Fig 4A, white bar in left panel). This approach allowed us to isolate the effect of V113L in its immediate historical context, while preserving all other substitutions separating AncNeoaves from AncPasserida, and thus enabling a focused test of its functional role in modulating Q111E. We confirmed that introducing the V113L substitution reversed the effect of Q111E, changing its impact from increasing to decreasing CTS resistance (Fig. 4A, left panel; Table S4). This significant functional alteration between AncNeoaves and AncPasserida demonstrates the crucial role of V113L in the evolutionary transition to the modern grosbeak genotype. The fixation of V113L seems to have restricted the subsequent evolution of Q111E, potentially necessitating the fixation of the additional five substitutions in the modern grosbeak. This underscores how genetic context and epistatic interactions can shape the accessibility of adaptive mutations, and how mutations that are initially deleterious can become beneficial again through subsequent permissive or compensatory changes.

To test how the functional effects of 6subs changed over time, we introduced them into the three reconstructed ancestral backgrounds. On the most ancient background, AncAves, 6subs had no significant effect on CTS resistance (Fig 4A, middle panel). On AncNeoaves, they slightly decreased resistance—an effect that was slightly stronger than the resistance-decreasing effect of Q111E alone on AncPasserida. In contrast, on AncPasserida, 6subs produced a significant increase in resistance (Fig 4A, middle panel). Thus, despite V113L reversing Q111E’s effect in AncPasserida, 6subs still conferred CTS resistance with or without Q111E. When Q111E and 6subs were introduced together, they consistently produced a significant increase in CTS resistance across all ancestral backgrounds (Fig. 4A, right panel; Table S4).

These findings illustrate how epistatic interactions shaped the emergence of resistance: both Q111E and 6subs could contribute to CTS resistance, but their effects varied with genetic context. This suggests a distinctive form of historical contingency in the protein’s evolution, in which intramolecular epistasis locks in the effects of certain beneficial mutations by making their functional impact dependent on the presence of earlier substitutions. Such interactions can open otherwise inaccessible evolutionary pathways, enabling the gradual assembly of complex, adaptive phenotypes like CTS resistance.

To further understand the evolutionary dynamics of these substitutions, we investigated whether there were functional trade-offs between toxin resistance and protein activity, which could have influenced the fixation of the modern grosbeak substitutions. Across all ancestral backgrounds, we found that Q111E has a minor negative effect on protein activity (Fig. 4B left panel; Table S4). In contrast, 6subs had consistently positive effects on protein activity when introduced on ancestral backgrounds, with significant effects on the AncPasserida and AncAves backgrounds (Fig. 4B, middle panel; Table S4). These results suggest that the evolution of CTS resistance via 6subs alone would have likely led to an increase in protein activity. When combined, Q111E and 6subs produced a mix of positive and negative effects on protein activity depending on the genetic background (Fig. 4B, right panel; Table S4). The combination had a neutral to negative effect on protein activity in AncAves and AncNeoaves, but significantly increased activity in the more recent AncPasserida background (Table S4; Fig. 4B right panel; Table S4). This positive effect of the combined substitutions on activity in AncPasserida aligns with the negative effect of reverting these mutations in the modern grosbeak. Our results suggest that the epistatic interaction between Q111E and 6subs balances the trade-off between CTS resistance and native protein activity, providing further evidence that intramolecular epistasis facilitated the evolutionary assembly of the resistant genotype in the grosbeak. We confirmed that functional inferences were robust to statistical uncertainties in ancestral reconstruction (Table S6).

Our functional experiments established that the evolution of target-site insensitivity in the modern grosbeak Na,K-ATPase involved nonadditive interactions between Q111E and 6subs. To assess evidence for interactions between Q111E and individual substitutions comprising 6subs in grosbeaks, we investigated phylogenetic patterns of co-variation in amino acid states across the Na,K-ATPases orthologs of class Aves. We built a phylogeny of full-length ATP1A1 sequences of 360 birds and created a comprehensive catalogue of amino acid variation among birds (Fig. 5). Notably, Q111E has evolved independently at least seven times across distinct bird lineages, including the *Tinamiformes* (tinamous), *Cuculiformes* (cuckoos), *Psittaciformes* (parrots), *Trogoniformes* (trogons), *Galliformes* (game birds), and *Musophagiformes* (turacos). The grosbeak is the only songbird (*Passeriformes*) so far identified to have this substitution [33].

**Figure 5.**
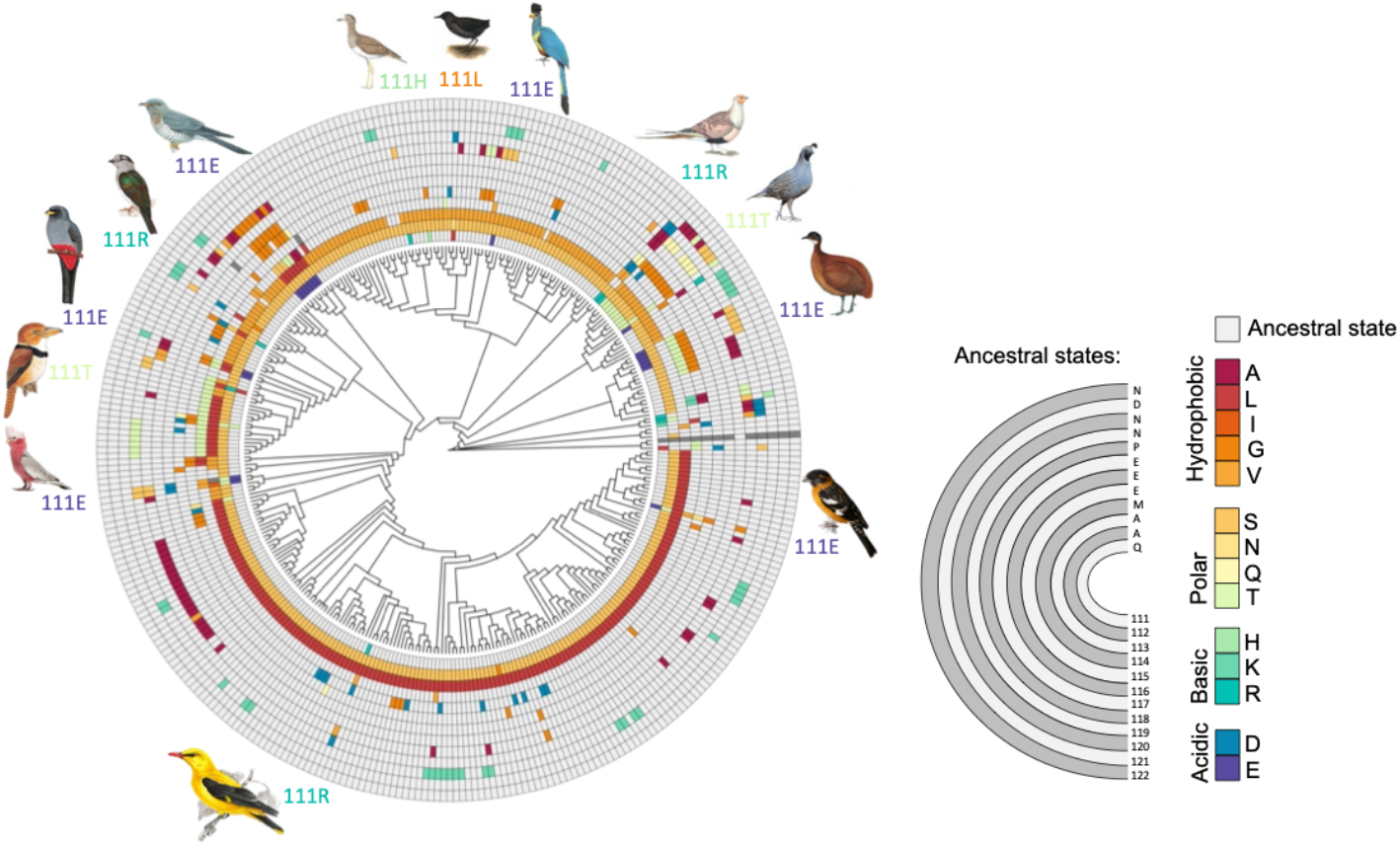
Circular phylogeny of birds showing amino acid states of the H1–H2 loop of the Na,K-ATPase ATP1A1 gene. Mutations within the H1–H2 loop have been associated with target-site insensitivity of the enzyme towards cardiotonic steroids, especially at sites 111 and 122. Notable substitutions are illustrated around the phylogeny.

Given that the avian ATP1A1 sequence is extremely well conserved across all species (mean branch length of 0.0037 substitutions per site), we investigated whether the Q111E substitution consistently co-occurs with other (potentially compensatory) substitutions in different lineages. Site 114, which is among the six substitutions comprising 6subs, was one of the eight sites whose substitution patterns fall within the top 5% for coevolutionary signal with Q111E (Fig. 5; Fig. S2). The substitution M114V in the grosbeak has evolved independently in a cockatoo clade where Q111E is also fixed. Further, all the closest relatives of the grosbeak that are known to be sensitive to the toxin retain the ancestral state at site 114 (Fig. 1; Fig. 5).

We expanded our coevolution analysis to identify additional sites that could be contributing to the evolution of target-site insensitivity in the grosbeak. We relaxed the coevolution analyses to identify other sites with substitutions that show a non-random co-occurrence with changes at position 111, regardless of whether that change is to glutamate (E) or another amino acid. This leverages the fact that site 111, which is involved in the convergent evolution of target-site insensitivity across vertebrates [16], has repeatedly experienced substitutions across the avian phylogeny (at least 13 times; Fig. 5). Among the 5% sites with strongest coevolution signals with site 111, two sites – 112 and 116 – are part of 6subs (Fig. S3). The substitution S112T in the grosbeak, for example, occurs independently in the same cockatoo clade that carries M114V, while all the closest relatives of the grosbeak maintain the ancestral state at site 112. These results suggest that epistatic interactions between Q111E, S112T, M114V, and E116G also contribute to the evolution of target-site insensitivity in the grosbeak.

The conditional effects of multiple substitutions in the grosbeak Na,K-ATPase represent a unique evolutionary outcome within this broader pattern of repeated adaptation [16, 20, 23, 24, 28, 34]. In contrast, other species such as horned frogs (*Ceratophrys*) and garter snakes (*Thamnophis*), both of which prey on CTS-containing toads, have independently evolved alternative resistance mechanisms involving a single amino acid substitution [16]. Similarly, the yellow-throated sandgrouse (*Pterocles gutturalis*) has developed a distinct single-amino acid insertion in the H1–H2 extracellular loop, a modification also associated with CTS resistance in pyrgomorphid grasshoppers [35-37]. These examples emphasise how different species have achieved the same functional outcome – resistance to CTS – through distinct molecular solutions, highlighting the diversity of adaptive strategies across species and trophic levels, and the complexity of evolutionary pathways to the convergent evolution of toxin resistance.

## Resource Availability

### Lead Contact

Further information and requests on methods can be directed to Dr. Hannah M. Rowland and Dr. Shabnam Mohammadi.

### Materials Availability

Plasmids used in this study have been deposited in Addgene (see Key Resources Table for names and numbers). This study did not generate new unique reagents.

### Data and Code

Data and code generated during this study are available through links provided in the Key Resources Table. Sequence data for ATP1A1 and ATP1B1 were obtained from genome assemblies under BioProject PRJNA545868 [38].

## Acknowledgements

We thank Guojie Zhang and Josefin Stiller for initial discussions and subsequent access to the genomic resources under BioProject PRJNA545868 [38]. We thank J. Stiller, M. Winters, K. Rohlfing, M. Röpcke, and K. Otto for critical reading of drafts of this manuscript. We thank V. Wagschal, C. Plate, F. Miazzi, and K. Barthold for assistance in the laboratory. This research was funded by the Max Planck Society to HMR.

## Author Contributions

**Conceptualisation:** SM, JFS, HMR; **Methodology:** SM, FGH, SHA, AE, SD, JFS, HMR; **Software**: SM, FGH, SHA, YD, HMR; **Validation:** SP; **Formal analysis:** SM, SP, FGH, SHA, YD, HMR; **Investigation:** SM, FGH, SHA, YD; **Resources:** FGH, SD; **Data Curation**: SM, SP, FGH, AE, HMR; **Writing - Original Draft**: SM, HMR; **Writing - Review & Editing:** SP, FGH, SHA, YD, AE, SD, JFS; **Visualization:** SM

## Key Resources Table

| Reagent or Resource | Source | Identifier |
| --- | --- | --- |
| **Bacterial and Virus Strains** |  |  |
| *Escherichia coli* MAX Efficiency™ DH10Bac Competent Cells | Thermo Fisher Scientific | Cat#10361012 |
| *Escherichia coli* XL 10-Gold Competent Cells | Agilent Technologies, La Jolla, CA, USA | Cat#200314 |
| **Experimental Models: Cell Lines** |  |  |
| Insect: Sf9 cells in Sf-900™ II SFM | Thermo Fisher | Cat#11496015 |
| **Technical devices** |  |  |
| Ultrasonic homogeniser Sonopuls 2070 | Bandelin Electronic Company, Berlin, Germany |  |
| Centrifuge 5840 R | Eppendorf AG, Hamburg, Germany |  |
| Ultra-Centrifuge L-80 | Beckmann Coulter GmbH, Krefeld, Germany |  |
| Freeze dryer Alpha 1-2 LD plus | Martin Christ Gefriertrocknungsanlagen GmbH, Osterode am Harz, Germany |  |
| Ultrasonic homogeniser Omni Sonic Ruptor 400 | Omni International Inc., Kennesaw, GA, USA |  |
| Cuvette spectrophotometer Ultrospec 2100 pro | Amersham Biosciences Europe GmbH, Freiburg im Breisgau, Germany |  |
| Microplate shaker BioShake iQ | Quantifoil Instruments, Jena, Germany |  |
| Microplate absorbance reader Bio-Rad Model 680 Bio-Rad iMark (spectrophotometers and software package) | **Bio-Rad Laboratories GmbH**, Feldkirchen, Germany |  |
| 96-well polystyrene flat-well plate | SARSTEDT AG & Co. KG, Nümbrecht, Germany | Cat#82.1581 |
| **Chemicals, Peptides, and Recombinant Proteins** |  |  |
| Insect-Xpress medium | Lonza, Walkersville, MD, USA | Cat#BE12-730P10 |
| Gentamycin | Roth, Karlsruhe, Germany | Cat#0233.1 |
| PEI MAX | Polysciences Inc., Warrington, PA, USA | CAS#49553-93-7 |
| HPLC grade water ROTIPURAN p.a., ACS water | Roth, Karlsruhe, Germany | Cat#HN68.2 |
| Bovine serum albumin (BSA) | Roth, Karlsruhe, Germany | CAS#90604-29-8 |
| Nitrocellulose membrane | Roth, Karlsruhe, Germany | Cat#HP42.1 |
| BlueBlock PF | SERVA Electrophoresis GmbH, Heidelberg, Germany | Cat#42591.01 |
| 4-Chloro-1-naphthol | Merck KGaA, Darmstadt, Germany | Cat#C8890 |
| Tween 20 | Merck KGaA, Darmstadt, Germany | CAS#9005-64-5 |
| 3,3',5,5'-tetramethylbenzidine (TMB) | Thermo Fisher Scientific Inc., Rockford, IL, USA | Cat#N301 |
| Parafilm | Merck KGaA, Darmstadt, Germany | Cat#P7793 |
| Ouabain octahydrate, 96% | Thermo Fisher Scientific Inc., Rockford, IL, USA | CAS#11018-89-6 |
| Adenosin-5-triphosphat Bis-(Tris)-salt hydrate (ATP) | Merck KGaA, Darmstadt, Germany | CAS#102047-34-7 |
| **Antibodies** |  |  |
| α5-antibody (primary antibody) | Developmental Studies Hybridoma Bank, University of Iowa, Iowa City, IA, USA | RRID: AB_2166869 |
| HRP-antibody (secondary antibody conjugated with horseradish peroxidase) | Dianova, Hamburg, Germany | RRID: AB_2617176 |
| **Deposited Data** |  |  |
| Raw data for recombinant Na,K-ATPase functional assays | This paper | Dryad doi: https://doi.org/ |
| ATP1A1 and ATP1B1 alignments and associated phylogenetic trees used to generate ancestral sequence reconstructions | This paper | Dryad doi: https://doi.org/ |
| R scripts and associated data tables used for statistical analyses and generating figures | This paper | Dryad doi: https://doi.org/ |
| **Recombinant DNA** |  |  |
| All plasmids generated in this study are listed in Table S1 | This paper | See Table S1 |
| **Software** |  |  |
| R | https://www.r-project.org/ | R version 4.1.1 |

## Methods

### Ancestral protein reconstruction

We compiled a dataset of avian ATP1A1 (n=370 species) and ATP1B1 (n=365 species) sequences from high-coverage genome assemblies (BioProject PRJNA545868 [38]) together with online sequences from nine reptiles [39-41]. Ancestral reconstruction was conducted on 343 ATP1A1 and 340 ATP1B1 sequences. Sequences were aligned using the MAFFT alignment algorithm [42], and ancestral reconstruction was estimated using the maximum likelihood (ML) approach implemented in IQ-TREE v 2.1.2 [43], with the aid of the Lazarus set of Python scripts for parsing results [44]. We constructed supertrees for avian sequences based on phylogenies corresponding to the most well-supported understanding of the species tree at the time of analysis [45-47]. Insertions were manually removed from marginal reconstructions, and ATP1A1 and ATP1B1 sequences of three ancestral NKAs in the grosbeak lineage were reconstructed and subsequently synthesised: AncAves, AncNeoaves, and AncPasserida. We also reconstructed and synthesised alternative ancestral reconstructions for altAncAves and altAncNeoaves by substituting sites with posterior probabilities below 0.8 when the ratio of the top two probabilities was less than 4:1. These alternative constructs were functionally assayed alongside the best estimates to confirm that minor uncertainty did not affect inferred function (Table S6). All sites in AncPasserida had posterior probabilities above 0.8, so no alternatives were generated.

### Construction of expression vectors

ATP1A1 and ATP1B1 genes of each recombinant NKA construct were synthesised (Invitrogen GeneArt) and codon optimised for *Spodoptera frugiperda*. The genes were cloned in pFastBac Dual (PFBD) plasmid with ATP1B1 under p10 promoter and ATP1A1 under P_PH_ promoter and confirmed by sequencing. The plasmid constructs were fully sequenced and deposited in the Addgene repository under accession numbers 196450 and 196454–70.

### Generation of recombinant viruses and transfection into Sf9 cells (*Spodoptera frugiperda*)

*Escherichia coli* DH10bac cells harboring the baculovirus genome (bacmid) and a transposition helper vector (Thermo Fisher Scientific; Cat#10361012) were transformed using a modified protocol of the manufacturer’s protocol with expression vectors containing the different gene constructs. Recombinant bacmids were selected using the blue-white screening method in agar plates containing kanamycin (50 μg/ml), gentamycin (7 μg/ml), tetracycline (10 μg/ml), X-gal (100 μg/ml) and IPTG (40 μg/ml). Positive (white) colonies were confirmed by PCR using the M13 primers (Forward primer: 5’-CCCAGTCACGACGTTGTAAAACG-3’, Reverse primer: 5’-AGCGGATAACAATTTCACACAGG-3’) as recommended by the manufacturer.

### Preparation of Sf9 membranes

Baculovirus-mediated transfection was performed using a modified protocol as described by Scholz and Suppmann [48]. Transfection mix was prepared in 100 μl PBS with 10 μg recombinant bacmid and 20 μl PEI-MAX (1mg/ml) per biological replicate. After 30 min of incubation, the transfection mix was added to actively growing Sf9 cells (8×106 cells/ml) in 10 ml of Insect-Xpress medium (Lonza; Cat#BE12-730P10) with 30 μg/ml gentamycin. Transfected cells were incubated at 27°C and 120rpm, and amplified viruses were harvested after 5 days by pelleting and discarding the cells (P0 virus stock). 500 μl of P0 virus stock was added to 5×10^6^ Sf9 cells/ml in 50ml Insect-Xpress media with 30 μg/ml gentamycin and cultured at 27°C and 120rpm in 250 ml conical flasks for 72h. Following incubation, Sf9 cells were harvested by centrifugation at 1650 × g for 10 min and the resulting cell pellets were immediately stored at –80°C.

### Membrane isolation from Sf9 cells

Frozen pellets were homogenised in 15ml homogenisation buffer (0.25 M sucrose, 2 mM ethylenediaminetetraacetic acid, and 25 mM HEPES/Tris; pH 7.0) on ice. Cell suspension was sonicated at 60 W (Sonopuls 2070; Bandelin Electronic Company, Berlin, Germany) for three 45 s intervals on ice. The cell suspension was then centrifuged for 30 min at 10,000 × g (J2-21 centrifuge, Beckmann-Coulter, Krefeld, Germany) at 4°C to remove debris. The supernatant was collected and further centrifuged for 60 min at 146,000 × g at 4°C (Ultra-Centrifuge L-80, Beckmann-Coulter) to pellet the cell membranes. The pelleted membranes were washed twice and resuspended in 1ml ROTIPURAN p.a., ACS water (Roth; Cat#HN68.2) and stored at −20 °C. Three biological replicates of each construct were used to conduct functional assays and protein concentrations were determined by ELISA.

### Protein Quanitification and Verification by ELISA and SDS-PAGE/western blotting

For the ELISA, we followed the protocol as described by Löptien et al.[49]. First, 100 µL of membrane isolates and relative standards were added to a 96-well polystyrene, slightly hydrophilic flat-well plate (Immulon 2 HB, Thermo Fisher Scientific, Massachusetts, USA; Cat#6302) in duplicate (two technical replicates). The standards were prepared from an aliquot of lyophilised *Pheucticus melanocephalus* membrane isolate as described by Löptien et al [49]. The plate was sealed and incubated overnight at 4°C (12–18 h). The next day, non-coated leftovers were removed from the wells and washed five times with washing buffer (Phosphate-buffered saline – 137 mM NaCl, 1.47 mM KH_2_PO_4_, 7.749 mM Na_2_HPO_4_, 2.683 mM KCl with 0.05 % Tween 20). Next, non-specific binding sites were blocked by adding 200 µL of blocking buffer (1% BSA in PBS + 0.02% Tween 20) and incubating at room temperature for 2 h. Following the blocking, the wells were washed another five times. NKAs were detected by adding 50 µL of primary antibody solution containing 2 µg/mL antibody (α5-antibody; Developmental Studies Hybridoma Bank, University of Iowa, Iowa City, IA, USA; RRID: AB_2166869) in blocking buffer incubating for 1 h at room temperature, then washed five times. To detect the primary antibody, 50 µL of secondary antibody solution containing 10 µg/mL HRP-conjugated goat-anti-mouse antibody (HRP-antibody; Dianova, Hamburg, Germany; RRID: AB_2617176) in blocking buffer was added and incubated for 1 h at room temperature. Following secondary antibody incubation, wells were washed seven times. The secondary antibody was stained by adding 100 µL 3,3’,5,5’ tetramethylbenzidine (TMB; Merck/Sigma-Aldrich; Cat#T4444) to each well, and then left to incubate at room temperature for 10 mins. Staining was stopped by adding 100 µL 0.5 M sulfuric acid to each well. The OD was read at 495 nm using a plate reader (BioRad Model 680 spectrophotometer and software package).

For the western blot, we followed the protocol described by Löptien et al.[49] In brief, 5 µL of protein were solubilised overnight in 8 µL 4 × Laemmli buffer (62.5 mM Tris HCl pH 6.8; 2 % SDS; 10 % glycerol; 5 % 2-mercaptoethanol; 0.001 % Bromophenol Blue) and 19 μL Millipore water, then separated on SDS gels containing 10% acrylamide. They were then blotted on nitrocellulose membrane (HP42.1, Roth). After blotting, the membrane was incubated with 1 × BlueBlock PF for 1 h. After blocking, the membranes were incubated at room temperature with the primary monoclonal antibody α5 (Developmental Studies Hybridoma Bank, University of Iowa, Iowa City, IA, USA) for 1.5 hours. Since only membrane proteins were isolated from transfected cells, detection of the α subunit also indicates the presence of the β subunit. The primary antibody was detected using a goat-anti-mouse secondary antibody conjugated with horseradish peroxidase (Dianova, Hamburg, Germany). The staining of the precipitated polypeptide-antibody complexes was performed by repetitively pipetting the detecting substrate (0.035 % H_2_O_2_, and 0.01 % 4-Chloro-1-naphthol in 0.05 M, pH 7.5 Tris HCl) over the membrane. See Figure S4.

### Ouabain inhibition assay

To determine the sensitivity of each NKA construct to the water-soluble cardiotonic steroid ouabain (Ouabain octahydrate 96%; Acrōs Organic; Cat#AC161730010s), 100 ug of each protein was pipetted into each well in a nine-well row on a 96-well flat-bottom microplate containing stabilising buffers (see buffer formulas in [34]. The first six wells in a nine-well row were exposed to exponentially decreasing concentrations of ouabain (10^−3^ M, 10^−4^ M, 10^−5^ M, 10^−6^ M, 10^−7^ M, 10^−8^ M, dissolved in distilled H_2_O). The seventh well was exposed to HPLC-grade water only (experimental control). The eighth well was exposed to an inhibition buffer lacking KCl and containing 10^− 2^M ouabain (ouabain octahydrate 96%; Acrōs Organic; Cat#AC161730010s), which completely inhibits the NKA’s ATPase activity and therby allows measurement of background ATPase activity (see Petschenka et al.[34]). The proteins were incubated at 37°C and 200 rpms for 10 minutes on a microplate shaker (BioShake iQ; Quantifoil Instruments, Jena, Germany; Cat#1808-0506). Next, ATP (Adenosin-5-triphosphat Bis-(Tris)-salt hydrate; Merck/Sigma-Aldrich; CAS#102047-34-7) was added to each well and the proteins were incubated again at 37°C and 200 rpm for 20 minutes. The activity of Na,K-ATPases following ouabain exposure was determined by quantification of inorganic phosphate (Pi) released from enzymatically hydrolysed ATP. Reaction Pi levels were measured according to the procedure described by Taussky and Shorr [50] (see Petschenka et al [34]). All assays were run in duplicate and the average of the two technical replicates was used for subsequent statistical analyses. Two constructs were run in Absorbance for each well and were measured at 650 nm with a plate absorbance reader (BioRad Model 680 spectrophotometer and software package).

### ATP hydrolysis assay

To determine the functional efficiency of different Na,K-ATPase constructs, we calculated the amount of Pi hydrolysed from ATP per mg of protein per minute. The measurements were obtained from the same assay as described above. In brief, absorbance from the experimental control reactions, in which 100 ug of protein was incubated without any inhibiting factors (i.e., ouabain or buffer excluding KCl), was measured and translated to mM Pi from a standard curve that was run in parallel (1.2 mM Pi, 1 mM Pi, 0.8 mM Pi, 0.6 mM Pi, 0.4 mM Pi, 0.2 mM Pi, 0 mM Pi). ATP hydrolysis measurements were always taken after the second freeze-thaw of each sample.

### Statistical analyses

Background phosphate absorbance levels from reactions with inhibiting factors were used to calibrate phosphate absorbance in wells measuring ouabain inhibition and in the control wells [34]. For ouabain sensitivity measurements, calibrated absorbance values were converted to percentage non-inhibited protein activity based on measurements from the control wells [34]. These data were plotted and log_10_ IC_50_ values were obtained for each biological replicate from nonlinear fitting using a four-parameter logistic curve, with the top asymptote set to 100 and the bottom asymptote set to zero. Curve fitting was performed with the nlsLM function of the minipack.lm library in R. For comparisons of protein activity, the calculated Pi concentrations of protein assayed in the absence of ouabain were converted to nmol Pi/mg protein/min. IC_50_ values were log_10_-transformed prior to analysis to better meet the assumptions of normality and homogeneity of variance. All statistical analyses were implemented in R. Data were plotted using the ggplot2 package in R.

We first analysed the effects of substitutions in the modern grosbeak NKA. We ran a 2-way ANOVA to assess the main effects of point mutations and their interaction, followed by planned contrasts to identify significant differences between recombinant proteins (Table S2). We then used linear regression to estimate the effect sizes of the explanatory variables and to assess their additive and nonadditive effects (Table S2). In a second set of analyses, we determined the background dependence of substitutions (i.e., interaction between genetic background and amino acid substitution) with respect to CTS resistance (log10 IC_50_) and ATPase activity (Table S5). Specifically, we tested whether the effects of a substitution X -> Y are equal on different backgrounds (null hypothesis: X->Y (background 1) = X->Y (background 2)). All statistical analyses were implemented in R. Data were plotted using the ggplot2 package in R.

### Identifying correlated substitutions

We used BayesTraits [51] to detect sites across the ATP1A1 bird phylogeny that exhibit correlated evolution with substitution Q111E or site 111, hereafter referred to as substitution and site models, respectively. Using the reconstructed ancestral sequence for the avian ATP1A1, we recoded each amino acid state among extant sequences of the multi-species alignment into ancestral ‘0’ and derived ‘1’ states and used these and the phylogeny with estimated branch lengths as inputs for BayesTraits. For the substitution model, site 111 was recoded as ‘1’ only for lineages that have 111E fixed; for the site model, site 111 was recoded as ‘1’ in every lineage with a substitution at this site (e.g., Q111X). These model specifications allowed us to use BayesTraits to fit a continuous-time Markov model to estimate transition rates between discrete, binary traits and estimate the best fitting model describing their joint evolution on a phylogeny. We excluded singleton sites and sites with more than 80% gaps, as these sites would be of little information. This resulted in an analysis of 144 variant sites.

With the substitution model, we tested whether the rate at which the substitution Q111E occurs was dependent on all other variant sites; with the site model, we tested whether the rate of evolution at site 111 was dependent on all other variant sites. Both coevolution models have a null independent and an alternative dependent model. The null independent model assumes that the two sites evolve independently, and the alternative dependent model assumes that the sites are correlated such that the change at one site is dependent on the state at the other site. Because the null model is a general form of the alternative model, both models can be compared under a likelihood ratio test (LRT) with degrees of freedom (df) equal to the difference in the number of parameters between models. Because the median branch length of the tree is 0.0019 substitutions per site, making multiple changes at a single site or reversions to the ancestral state unlikely, we set the rates of transition to the ancestral state to zero. After this restriction, the null independent model had four free transition parameters. To test for dependence, we imposed two additional restrictions to the model: one forcing the transition rate at site one to be fixed regardless of the state of site 2 (q13=q24), and a second forcing the transition rate at site 2 to be fixed regardless of the state of site 1 (q12=q34). This effectively tests whether the transition rate is affected by the state of either site leaving the model with only two free transition parameters (LRT with df=2). To run the analysis, the phylogeny branch lengths were scaled using BayesTraits to have a mean length of 0.1; to increase the chance of finding the true maximum likelihood, we set MLTries to 250.

