## Supplementary Materials for "Historical Contingency Shapes Toxin Resistance in a Specialist Avian Predator"

**Table S1.** List of gene constructs used to test functional effects of modern black-headed grosbeak ATP1A1 amino acid mutations on the modern Na,K-ATPase and on its ancestral variants. The corresponding ATP1B1 gene of each recombinant protein construct was co-expressed with ATP1A1. All grosbeak constructs contained the grosbeak ATP1B1 gene. All ancestral constructs contained the corresponding ancestral ATP1B1 gene.

| **Construct Name** | **Engineered substitution(s)** | **Description** | **Addgene ID** |
|---|---|---|---|
| Grosbeak | none | Grosbeak wildtype ATP1A1. | 196465 |
| Grosbeak+E111Q | E111Q | Grosbeak with site 111 reverted to ancestral state. | 196466 |
| Grosbeak+6sub | T112A<br>L113A<br>V114M<br>N115E<br>G116E<br>G118P | Grosbeak with sites 112-118 reverted to ancestral state. | 196467 |
| Grosbeak+0subs | E111Q<br>T112A<br>L113A<br>V114M<br>N115E<br>G116E<br>G118P | Grosbeak with entire H1-H2 loop reverted to ancestral state. | 218154 |
| AncPass | none | Ancestral Passerida ATP1A1. | 196461 |
| AncPass+Q111E | Q111E | Ancestral Passerida ATP1A1 with site 111 forward mutated to modern grosbeak state. | 196462 |
| AncPass+Q11E1+L113V | Q111E<br>L113V | Ancestral Passerida ATP1A1 with site 111 forward mutated to modern grosbeak state and with site 113 reverted to the ancestral Neoaves state. | 196470 |
| AncPass+6sub | A112T<br>A113L<br>M114V<br>E115N<br>E116G<br>P118G | Ancestral Passerida ATP1A1 with sites 112-118 forward mutated to modern grosbeak state. | 196463 |
| AncPass+7sub | Q111E<br>A112T<br>A113L<br>M114V<br>E115N<br>E116G<br>P118G | Ancestral Passerida ATP1A1 with the entire H1-H2 loop forward mutated to the modern grosbeak state. | 196464 |
| AncNeo | none | Ancestral Neoaves ATP1A1. | 196457 |
| AncNeo+Q111E | Q111E | Ancestral Neoaves ATP1A1 with site 111 forward mutated to modern grosbeak state. | 196458 |
| AncNeo+6sub | A112T<br>A113L<br>M114V<br>E115N | Ancestral Neoaves ATP1A1 with sites 112-118 forward mutated to modern grosbeak state. | 196459 |

| | | | |
|---|---|---|---|
| | E116G<br>P118G | | |
| AncNeo+7sub | Q111E<br>A112T<br>A113L<br>M114V<br>E115N<br>E116G<br>P118G | Ancestral Neoaves ATP1A1 with the entire H1-H2 loop forward mutated to the modern grosbeak state. | 196460 |
| AncAves | none | Ancestral Aves ATP1A1. | 196450 |
| AncBird+Q111E | Q111E | Ancestral Aves ATP1A1 with sites 111 forward mutated to modern grosbeak state. | 196454 |
| AncAves+6sub | A112T<br>A113L<br>M114V<br>E115N<br>E116G<br>P118G | Ancestral Aves ATP1A1 with sites 112-118 forward mutated to modern grosbeak state. | 196455 |
| AncAves+7sub | Q111E<br>A112T<br>A113L<br>M114V<br>E115N<br>E116G<br>P118G | Ancestral Aves ATP1A1 with the entire H1-H2 loop forward mutated to the modern grosbeak state. | 196456 |
| *altAncAves | - | - | 196468 |
| *altAncNeo | - | - | 196469 |

*Two additional constructs (altAncBird and altAncNeo) were engineered with the next probable protein sequences of the AncBird and AncNeo constructs and were produced to assure our ancestral sequence reconstructions expressed robust proteins. The posterior probabilities for all amino acid sites of AncPass were above our threshold of 0.80 and thus no alternative protein was inferred for that construct.

**Table S2.** Statistical analysis of ouabain sensitivity ($IC_{50}$) and ATPase activity of recombinant modern black-headed grosbeak proteins. Significant p values are highlighted in bold. Levene's Test for homogeneity of variance for $IC_{50}$: $F_{3,8}$= 0.0572, p= 0.9808; and for ATPase: $F_{3,8}$= 0.6901, p= 0.5832.

| **(Explanatory Variables) ANOVA** | Ouabain sensitivity log10($IC_{50}$) | | | | ATPase activity nmol Pi/(mg protein*min) | | | |
|---|---|---|---|---|---|---|---|---|
| | df | MS | F | p value | df | MS | F | p value |
| E111 | 1 | 1.0489 | 40.521 | **0.000217** | 1 | 61.93 | 32.98 | **0.000433** |
| 6subs | 1 | 0.2387 | 9.222 | **0.016142** | 1 | 21.91 | 11.67 | **0.009146** |
| E111:6subs | 1 | 0.4205 | 16.246 | **0.003785** | 1 | 0 | 0 | 0.983803 |
| Residuals | 8 | 0.0259 | | | 8 | 1.88 | | |

| **(Explanatory Variables) Linear regression** | Ouabain sensitivity log10($IC_{50}$) | | | | ATPase activity nmol Pi/(mg protein*min) | | | |
|---|---|---|---|---|---|---|---|---|
| | Est | SE | t | p value | Est | SE | t | p value |
| Intercept | -6.0708 | 0.0929 | -65.355 | **3.34e-12** | 2.210 | 0.791 | 2.794 | **0.0234** |
| E111 | 0.2169 | 0.1314 | 1.651 | 0.13733 | 4.527 | 1.119 | 4.046 | **0.0037** |
| 6subs | -0.0923 | 0.1314 | -0.703 | 0.50212 | 2.686 | 1.119 | 2.400 | 0.0431 |
| E111:6subs | 0.7488 | 0.1858 | 4.031 | **0.00378** | 0.033 | 1.582 | 0.021 | 0.9838 |

| Dependent Variable | Contrast Comparison | *df* | *t* | *p* | *d* | 95% CI |
|---|---|---|---|---|---|---|
| $IC_{50}$ | **WT – (E111Q)** | **8** | **7.35** | **< .001*** ** | **6** | **[5.50, 6.19]** |
| | **WT – (6subs)** | **8** | **5** | **.001** ** | **4.08** | **[3.61, 4.41]** |
| | **WT – (E111Q-6subs)** | **8** | **6.65** | **< .001*** ** | **5.43** | **[4.79, 6.17]** |
| | **E111Q - 6subs** | **8** | **-2.35** | **.046*** | **-1.92** | **[-4.05, 0.25]** |
| | E111Q – (E111Q-6subs) | 8 | -0.7 | 0.502 | -0.57 | [-2.51, 0.73] |
| | 6subs – (E111Q-6subs) | 8 | 1.65 | 0.137 | 1.35 | [-0.73, 3.39] |
| ATPase activity | **WT – (E111Q)** | **8** | **4.08** | **.004** ** | **3.33** | **[2.54, 6.02]** |
| | **WT – (6subs)** | **8** | **2.43** | **.041*** | **1.98** | **[0.55, 3.43]** |
| | **WT – (E111Q-6subs)** | **8** | **6.48** | **< .001*** ** | **5.29** | **[4.44, 7.02]** |
| | E111Q – (6subs) | 8 | -1.65 | .138 | -1.34 | [-5.98, 1.54] |
| | **E111Q – (E111Q-6subs)** | **8** | **2.40** | **.043*** | **1.96** | **[0.84, 3.40]** |
| | **6sub – (E111Q-6subs)** | **8** | **4.05** | **.004** ** | **3.30** | **[1.95, 8.73]** |

**Table S3.** Amino acid substitutions across the length of the ATP1A1 protein. 111-122 represents the first extracellular loop that interacts with cardiotonic steroids

| | 11 | 15 | 17 | 18 | 19 | 20 | 21 | 33 | 53 | 58 | 72 | 83 | 102 | 109 | 111 | 112 | 113 | 114 | 115 | 116 | 117 | 118 | 119 | 120 | 121 | 122 | 127 | 135 | 157 | 165 |
|---|---|---|---|---|---|---|---|---|---|---|---|---|---|---|---|---|---|---|---|---|---|---|---|---|---|---|---|---|---|---|
| **Anc_Aves** | A | K | G | K | K | E | K | S | S | P | A | V | V | G | Q | A | A | M | E | E | E | P | N | N | D | N | V | I | M | I |
| **Anc_Neoaves** | . | . | . | . | . | . | R | S | . | S | T | . | I | . | . | S | V | . | . | . | . | . | . | . | . | . | . | . | . | V |
| **Anc_ Passerida** | G | . | . | . | . | . | R | V | . | S | . | . | I | . | . | S | L | . | . | . | . | . | . | . | . | . | I | . | L | V |
| **Grosbeak** | G | G | — | — | — | — | . | V | N | S | . | I | I | A | E | T | L | V | N | G | . | G | . | . | . | . | I | V | L | V |

| | 171 | 225 | 226 | 248 | 250 | 253 | 279 | 287 | 409 | 412 | 417 | 429 | 462 | 467 | 513 | 527 | 550 | 560 | 568 | 569 | 574 | 676 | 875 | 879 | 888 | 978 | 993 | 997 | 1012 |
|---|---|---|---|---|---|---|---|---|---|---|---|---|---|---|---|---|---|---|---|---|---|---|---|---|---|---|---|---|---|
| **Anc_Aves** | L | T | N | R | V | N | T | I | P | S | V | N | E | N | S | T | H | E | E | V | E | T | S | I | V | P | V | V | K |
| **Anc_Neoaves** | M | S | . | . | I | S | M | L | A | T | . | . | . | Y | . | M | . | . | . | . | D | I | G | . | I | L | L | I | . |
| **Anc_ Passerida** | M | S | H | . | I | S | M | L | A | T | I | G | Q | Y | T | M | . | D | D | L | D | L | G | . | I | L | . | . | R |
| **Grosbeak** | M | S | H | H | I | S | M | . | A | T | I | G | Q | Y | T | M | Y | . | D | L | D | L | G | L | I | L | . | . | . |

**Table S4.** Statistical analysis of ouabain sensitivity and ATPase activity of recombinant ancestral bird proteins. Significant p values are highlighted in bold. Levene's Test for homogeneity of variance for $IC_{50}$: $F_{11,42}=0.3578$, $p=0.9652$; and for ATPase: $F_{11,43}=0.5682$, $p=0.8433$.

| **(Explanatory Variables) ANOVA** | Ouabain sensitivity log10($IC_{50}$) | | | | ATPase activity nmol Pi/(mg protein*min) | | | |
|---|---|---|---|---|---|---|---|---|
| | df | MS | F | p value | df | MS | F | p value |
| Background | 2 | 0.02541 | 0.3566 | 0.70723 | 2 | 5.8227 | 8.7952 | **0.004448** |
| Sub (Q111E) | 1 | 0.50243 | 7.0519 | **0.02096** | 1 | 3.5194 | 5.3161 | **0.039787** |
| Background:Sub | 2 | 0.37986 | 5.3316 | **0.02204** | 2 | 0.7870 | 1.1888 | 0.338043 |
| Residuals | 12 | 0.07125 | | | 12 | 0.6620 | | |

| **(Explanatory Variables) ANOVA** | Ouabain sensitivity log10(IC50) | | | | ATPase activity nmol Pi/(mg protein*min) | | | |
|---|---|---|---|---|---|---|---|---|
| | df | MS | F | p value | df | MS | F | p value |
| Background | 2 | 0.96406 | 17.3867 | **0.0002852** | 2 | 74.556 | 97.408 | **3.816e-8** |
| Sub (6subs) | 1 | 0.10152 | 1.8309 | 0.2009592 | 1 | 192.568 | 251.592 | **2.049e-9** |
| Background:Sub | 2 | 0.33255 | 5.9975 | **0.0156447** | 2 | 40.317 | 52.674 | **1.143e-6** |
| Residuals | 12 | 0.05545 | | | 12 | 0.765 | | |

| **(Explanatory Variables) ANOVA** | Ouabain sensitivity log10(IC50) | | | | ATPase activity nmol Pi/(mg protein*min) | | | |
|---|---|---|---|---|---|---|---|---|
| | df | MS | F | p value | df | MS | F | p value |
| Background | 2 | 0.17773 | 3.3059 | 0.0718400 | 2 | 9.8680 | 8.8145 | **0.004413** |
| Sub (Q111E+6subs) | 1 | 1.43684 | 26.7254 | **0.0002333** | 1 | 0.5243 | 0.4683 | 0.506766 |
| Background:Sub | 2 | 0.00603 | 0.1122 | 0.8947943 | 2 | 3.1662 | 2.8282 | 0.098557 |
| Residuals | 12 | 0.05376 | | | 12 | 1.1195 | | |

**Table S5.** Contrast comparisons of ouabain sensitivity (IC50) and ATPase activity between different recombinant ancestral bird proteins. Significant p values are highlighted in bold and denoted by asterisks. Relevant comparisons, i.e., between wildtype proteins (WT) and their mutated counterparts (Q111E, 6subs, and Q111E+6subs), are highlighted in grey. Significance codes: 0 '***'; 0.001 '**'; 0.01 '*'.

| Dependent Variable | Comparison | *df* | *t* | *p* | *d* | 95% CI |
|---|---|---|---|---|---|---|
| AncAves IC50 | **WT – (Q111E)** | **8** | **-5.05** | **.001*****| **-4.12** | **[-5.44, -3.08]** |
| | WT – (6subs) | 8 | -0.71 | .500 | -0.58 | [-2.07, 3.34] |
| | **WT – (Q111E+6subs)** | **8** | **-4.09** | **.003**** | **-3.34** | **[-5.36, -2.02]** |
| | **Q111E – (6subs)** | **8** | **4.34** | **.002**** | **3.54** | **[3.07, 4.07]** |
| | Q111E – (Q111E+6subs) | 8 | 0.95 | .368 | 0.78 | [-1.65, 4.51] |
| | 6subs – (Q111E+6subs) | **8** | **-3.39** | **.010**** | **-2.76** | **[-3.78, -2.03]** |
| AncAves ATPase | WT – (Q111E) | 8 | 2.19 | .060 | 1.79 | [-0.12, 3.86] |
|  | **WT – (subs6)** | **8** | **-12.49** | **< .001***** | **-10.19** | **[-12.01, -8.39]** |
| | WT – (Q111E+6subs) | 8 | 1.23 | .255 | 1.00 | [-1.10, 2.86] |
|  | **Q111E – (subs6)** | **8** | **-14.68** | **< .001***** | **-11.98** | **[-14.49, -9.95]** |
| | Q111E – (Q111E+6subs) | 8 | -0.97 | .363 | -0.79 | [-2.10, 1.20] |
|  | **subs6 – (Q111E+6subs)** | **8** | **13.71** | **< .001***** | **11.20** | **[9.66, 13.25]** |
| AncNeo IC50 | WT – (Q111E) | 8 | -1.78 | .113 | -1.45 | [-2.66, -0.18] |
| | WT – (6subs) | 8 | 1.26 | .243 | 1.03 | [-1.45, 3.00] |
| | WT – (Q111E+6subs) | 8 | -2.21 | .058 | -1.80 | [-3.16, -0.21] |
| | **Q111E – (subs6)** | **8** | **3.04** | **.016*** | **2.48** | **[0.70, 4.66]** |
| | Q111E – (Q111E+6subs) | 8 | -0.43 | .679 | -0.35 | [-2.03, 2.94] |
| | **subs6 – (Q111E+6subs)** | **8** | **-3.47** | **.008**** | **-2.83** | **[-4.71, -0.75]** |
| AncNeo ATPase | WT – (Q111E) | 8 | 0.79 | .454 | 0.64 | [-2.80, 3.52] |
| | WT – (6subs) | 8 | -0.96 | .367 | -0.78 | [-4.20, 0.85] |
| | **WT – (Q111E+6subs)** | **8** | **2.34** | **.047*** | **1.91** | **[-0.55, 5.48]** |
| | Q111E – (6subs) | 8 | -1.74 | .119 | -1.42 | [-2.78, 0.21] |
| | Q111E – (Q111E+6subs) | 8 | 1.56 | .158 | 1.27 | [-0.48, 2.90] |
| | **6subs – (Q111E+6subs)** | **8** | **3.30** | **.011*** | **2.69** | **[1.34, 4.11]** |
| AncPass IC50 | WT – (Q111E) | 8 | 1.02 | .336 | 0.84 | [-1.37, 2.78] |
| | ***WT – (Q111E(+L113V))*** | **8** | **-2.69** | **.028*** | **-2.19** | **[-4.66, 0.87]** |
| | **WT – (6subs)** | **8** | **-3.16** | **.013*** | **-2.58** | **[-4.36, -1.12]** |
| | **WT- (Q111E+6subs)** | **8** | **-2.62** | **.030*** | **-2.14** | **[-4.33, 0.59]** |
| | **Q111E – (6subs)** | **8** | **-4.18** | **.003**** | **-3.41** | **[-4.06, -2.62]** |
| | **Q111E – (Q111E+6subs)** | **8** | **-3.65** | **.007**** | **-2.98** | **[-4.36, -1.65]** |
| | 6subs – (Q111E+6subs) | 8 | 0.53 | .609 | 0.43 | [-1.09, 2.27] |
| AncPass ATPase | WT – (Q111E) | 8 | 0.54 | .607 | 0.44 | [-1.22, 1.65] |
| | *WT – (Q111E(+L113V))* | 8 | -0.03 | .977 | -0.02 | [-1.11, 1.63] |
|  | **WT – (6subs)** | **8** | **-10.57** | **< .001***** | **-8.63** | **[-8.82, -8.21]** |
| | WT – (Q111E+6subs) | 8 | -1.51 | .171 | -1.23 | [-6.15, 3.56] |
|  | **Q111E – (6subs)** | **8** | **-11.10** | **< .001***** | **-9.07** | **[-9.47, -8.53]** |
| | Q111E – (Q111E+6subs) | 8 | -2.04 | .076 | -1.67 | [-8.35, 2.32] |
|  | **6subs – (Q111E+6subs)** | **8** | **9.06** | **< .001***** | **7.40** | **[7.09, 7.55]** |

Note – Contrast analyses were run twice with two versions of the ancestral Passerida WT vs. Q111E comparison. One dataset included the original ancPasserida+Q111E construct, which did not carry the additional L1113V substitution, and the other dataset included the ancPasserida+Q111E construct with the additional substitution.

**Table S6.** Results of Welch's t-test comparison between best and next-best ancestral sequence estimates for Anc_Aves and Anc_Neoaves.

| Comparison | Variable | t | df | p_value | CI_lower | CI_upper | mean_Anc | mean_altAnc |
|---|---|---|---|---|---|---|---|---|
| Anc_Aves vs altAnc_Aves | IC50 | 1.22043440 | 3.976660 | 0.2896901 | -0.2490968 | 0.6382374 | -6.206593 | -6.401163 |
| Anc_Aves vs altAnc_Aves | ATPase | 0.06085616 | 3.792780 | 0.9545443 | -2.4057588 | 2.5112210 | 5.934986 | 5.882255 |
| Anc_Neo vs altAnc_Neo | IC50 | 0.12424841 | 2.276140 | 0.9112397 | -1.7262322 | 1.8417022 | -6.109394 | -6.167129 |
| Anc_Neo vs altAnc_Neo | ATPase | -2.07420545 | 3.993673 | 0.1068310 | -3.8909393 | 0.5644903 | 3.581857 | 5.245081 |

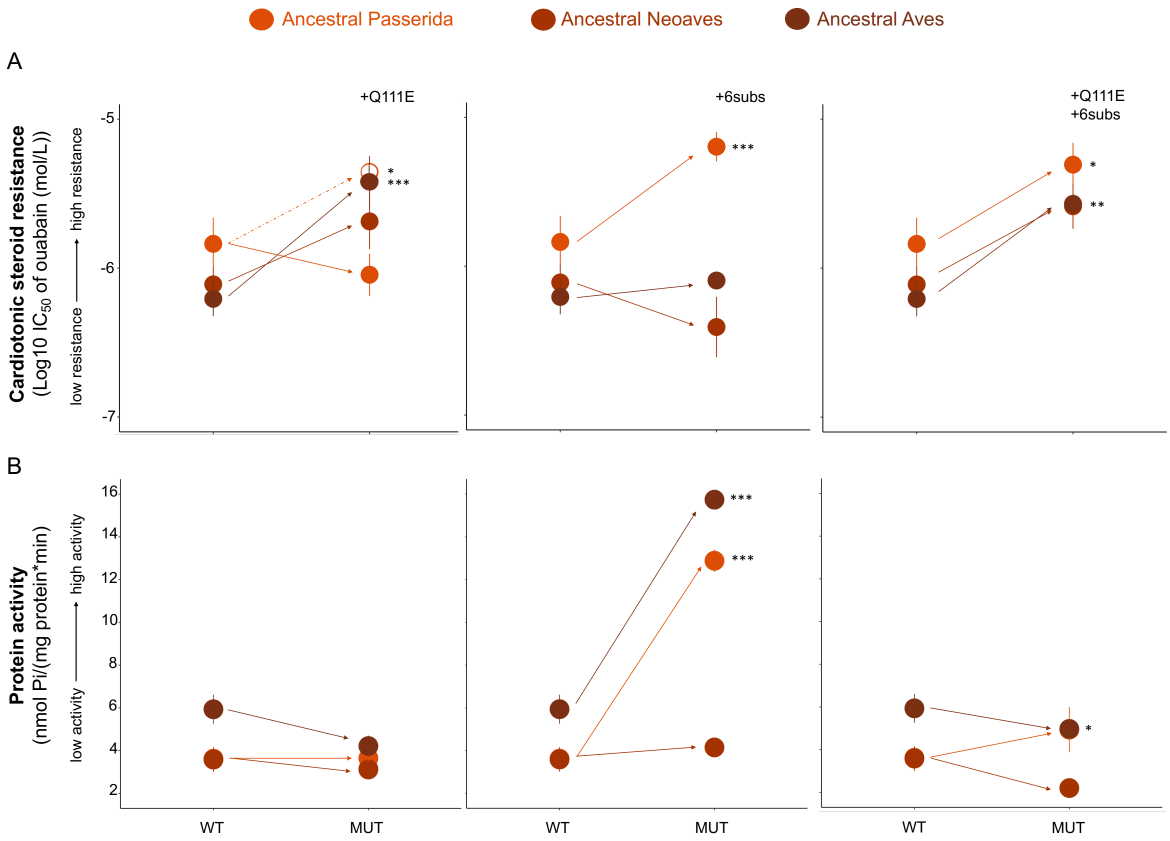


**Figure S1.** Functional properties of engineered Na,K-ATPases (NKAs) measuring the effects of modern black-headed grosbeak substitutions on increasingly more ancient genetic backgrounds (from orange to maroon). The graphs in panels A and B show the functional properties of wildtype ancestral NKAs (WT) in comparison to their mutagenized counterpart (MUT). Each graph (from left to right) shows the effects of one of three mutation variants (Q111E, 6subs, and Q111E+6subs). In panel A, a measure of CTS resistance (i.e., mean $\log_{10}IC_{50}$ ± SEM from three biological replicates) is plotted on the y axis. In panel B, a measure of protein activity (i.e., mean ATP hydrolysis rate ± SEM from three biological replicates) for the same proteins is plotted on the y axis. Significant differences (contrast analysis) between WT and MUT proteins are indicated by asterisks. In the Q111E graphs are an additional mutagenized construct of ancestral Passerida, engineered to carry Q111E as well as L113V. This additional construct is illustrated by an open circle and dashed arrow line. In panel B, this additional construct overlaps completely with the original and is therefore not visible. Significance codes: 0 '***'; 0.001 '**'; 0.01 '*'.

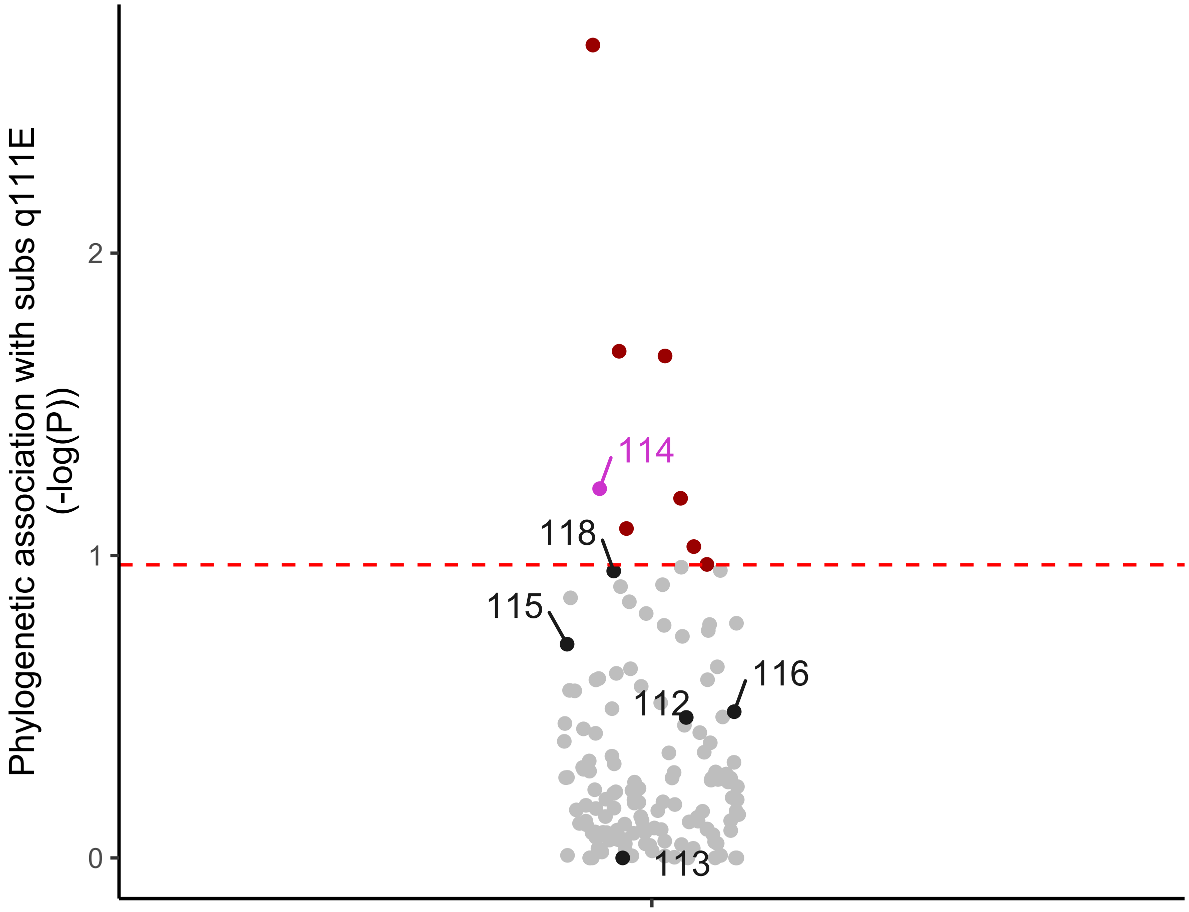


**Figure S2.** Distribution of p-values (phylogenetic association) with site Q111E.

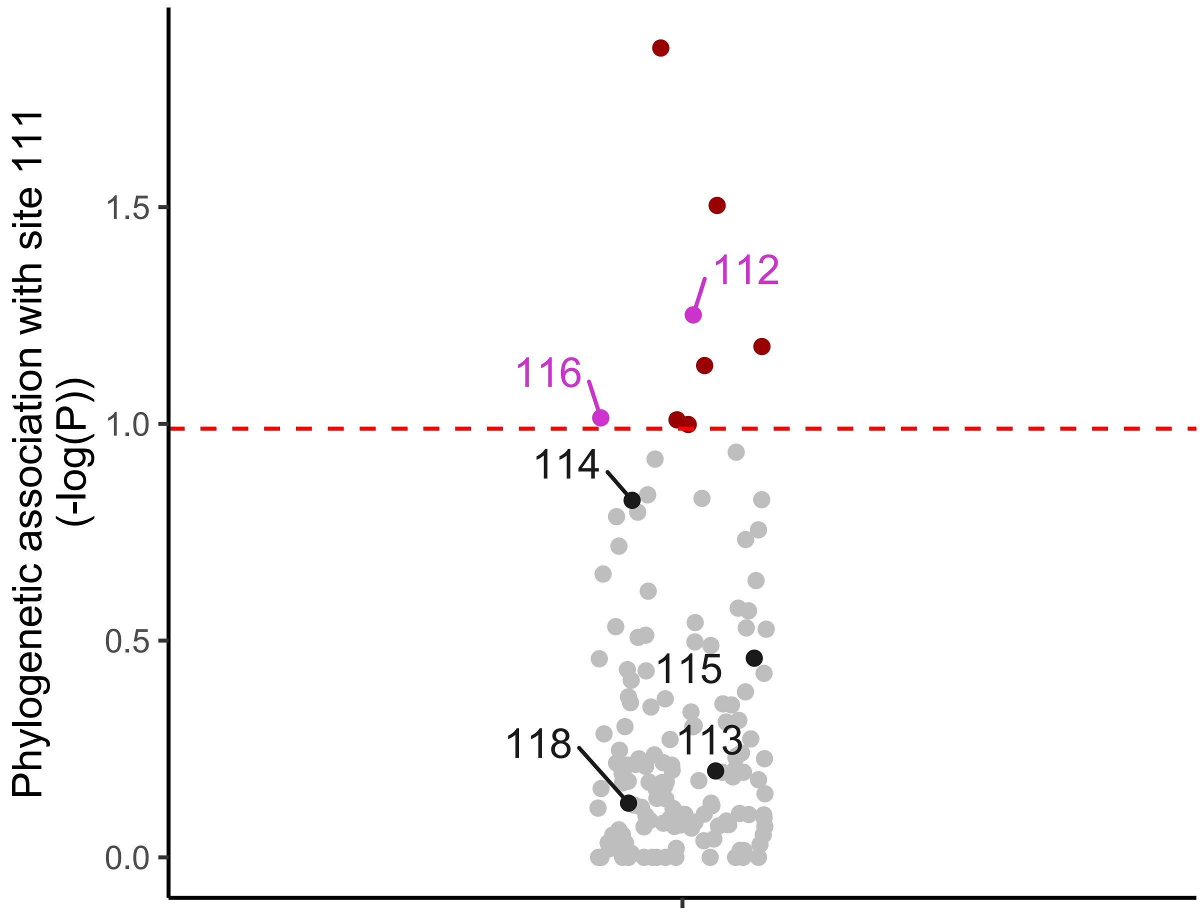


**Figure S3.** Distribution of p-values (phylogenetic association) with changes at site 111.

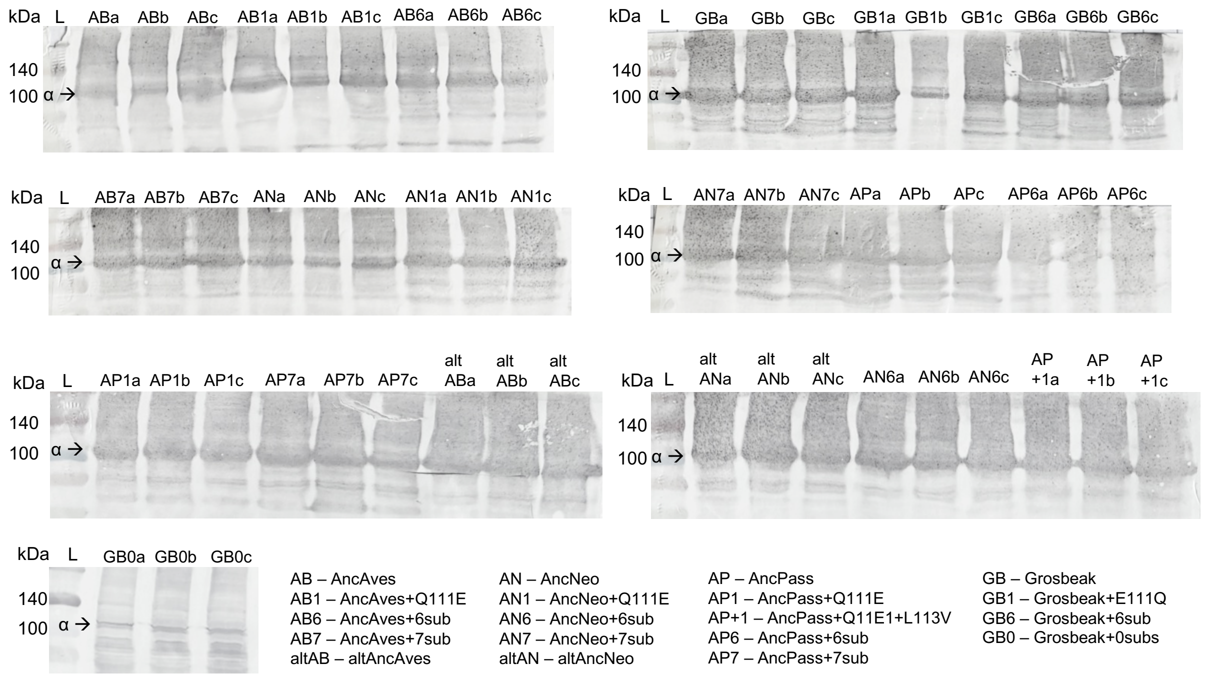


**Figure S4.** Western blot analysis of Na,K-ATPase with engineered ATP1A1 (α-subunits) produced in this study. The 110 kDa α-subunits are stained with the α5 monoclonal antibody followed by a horseradish peroxidase conjugated goat antimouse antibody. Samples represents three biological replicates of 19 different recombinant Na,K-ATPases (Table S1) produced through cell culture. Each scanned photo represents one western blot gel. The lanes are labelled according to the sample they contained, and the letter "L" indicates the protein ladder. For each western blot, 5 µl of membrane-isolated protein sample was used.
